# Design and fabrication of a low-cost microfluidic cartridge with integrated pressure-driven check valve for molecular diagnostics platforms

**DOI:** 10.1101/2022.12.29.522222

**Authors:** R. Scott Downen, Quan Dong, Julius Lee Chen, Zhenyu Li

**Affiliations:** Department of Biomedical Engineering, The George Washington University, Washington, D.C.

## Abstract

This paper describes the design, fabrication, and preliminary testing of a low-cost, easy to manufacture microfluidics cartridge capable of fluid storage and manipulation through a custom pressure-driven check valve. Cartridge components are fabricated using a desktop CNC and laser cutter, the check valve is fabricated using PDMS in a custom acrylic mold, and the components are assembled using a thermal diffusion welder. Following assembly, preliminary testing of the cartridge, including fluid manipulation and use for molecular diagnostics, was performed. To pull a sample into the lysing chamber, a vacuum over 1.4PSI was required. No opening of the valve to the reaction chamber was observed. Moving fluid across the custom valve from the lysing chamber to the reaction chamber then required a vacuum over 4.5PSI. Finally, a proof-of-concept demonstration of one potential application was performed using a custom benchtop LAMP system for molecular diagnostic testing. The low-cost nature of the design, ease of manufacturing, fluid storage and manipulation demonstrated make this design ideal for research and high-volume testing in low resource environments.

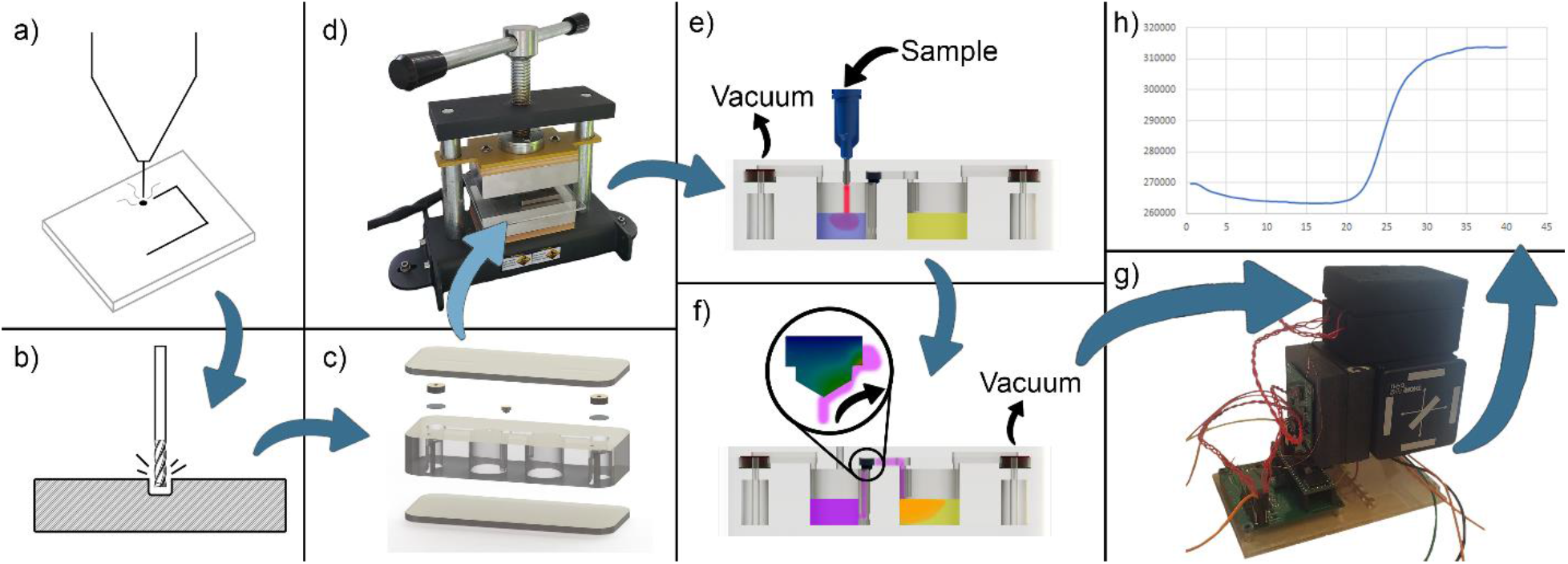

**Graphical abstract:** Custom cartridge is fabricated using a combination of a) benchtop laser cutter and b) benchtop micromilling machine. c) Components are then assembled with a 0.22µm micron filter and custom plug-style check valve. d) The cartridge assembly is then bonded using a thermal diffusion bonder. e) By pulling a vacuum through the first air trap, a sample can be pulled into the pre-filled lysing chamber. f) Pulling a vacuum through the second air trap, the lysed sample can then be pulled into the pre-filled reaction chamber. g) For a proof of concept, the filled cartridge was then tested in a custom benchtop Loop Mediated Isothermal System. Following a heating cycle, reaction fluoresce can be monitored. h) An S-Curve observed through the custom LAMP system, thus demonstrating feasibility of the cartridge for use with molecular diagnostic platforms.

## 1. Introduction

It can be argued that the importance of rapidly deployable, easy-to-use, and fast time-to-result genetic testing has never been more visible than during the COVID-19 pandemic. Meanwhile, it is estimated that the molecular diagnostics point of care market will grow to USD $4.84B by the year 2027 [1]. Additional techniques and process simplifications are necessary to further grow the field of point of care genetic testing, specifically targeting weaknesses which exist in a majority of commercially available systems such as high per-test cost, high prototype cost, and manual fluid manipulation requiring highly trained personnel. To develop these techniques in a time-efficient manner, availability of low-cost, easy-to-manufacture cartridges compatible with benchtop and portable molecular diagnostics platforms gain increased importance.

In low resource settings and prototype-based research laboratories, microfluidics-based diagnostics platforms offer several benefits over traditional, large fluid-volume systems including: low-cost testing due to reduced reagent volume needed per test; energy efficiency, as lower volumes of fluid may be heated more rapidly; sample-to-answer capabilities based on complex functions integrated in a single device; relatively fast output based on reduced fluid transport times within the system; and high portability for in-field testing and distribution for trials [2].

In a traditional laboratory environment, fabrication of fully integrated microfluidic devices capable of fluid manipulation and storage requires complex, large, and costly fabrication equipment. Standard high-volume processes such as injection molding [4], hot embossing [5], and thermoforming require master molds to transfer patterns, which typically makes it difficult to change designs rapidly, and multi-layer microfluidic patterns with these methods are often costly and time-consuming [4][6][7]. To eliminate this hurdle, CO_2_ laser machines have been adopted in many laboratories to create patterns in thermoplastics directly, bypassing the need for master molds [8]–[17]. Less frequently used, benchtop micromilling machines have also been demonstrated to provide adequate microfluidic fabrication capabilities for biomedical applications [18]–[23]. These systems are low-cost and able to provide fast results, making them a good candidate for rapid prototyping of microfluidic devices for rapid development, testing and validation [19].

By combining laser machining and micromilling techniques, benefits from each process can be harnessed for rapid, low-cost fabrication of cartridges adequate for optically based molecular diagnostics platforms. Using these techniques, a one-way check valve mold may also be fabricated to allow chamber separation and controlled fluid movement.

As discussed by Liu et al., microvalves are critical to the success of microfluidics-based molecular diagnostics platforms and have several requirements. First, for high temperature applications a great deal of pressure can be generated due to degassing and air expansion. If the valve fails, the fluid will move into an unwanted chamber, resulting in a failed test [3]. Second, as the valve will come in contact with a reagent solution, it must not interfere or inhibit the reaction. Lastly, the valve must open easily to allow fluid flow when desired [3]. With these requirements, the ability to quickly adapt a microvalve to changing cartridge parameters becomes critical to the success of the cartridge overall.

For fluid manipulation, check valve pumps, the most common pump type at a macroscale, were the first concept to be miniaturized [28]. Traditional check valve pumps contain an actuator, a pump membrane, a dead-volume pump chamber, and two check valves operated by pressure. Many check valve designs have been proposed for microfluidic systems, including ring mesa, cantilever, disc, V-shape, membrane, and floating type [29]. The present work features a one-way plug type check valve.

Using similar rapid prototyping methods, several groups have worked towards developing microfluidic cartridges with integrated actuators, microvalves, and micropumps [24]–[27]. However, many of these mechanisms of fluid manipulation require more expensive control systems and complicated manufacturing techniques, thus reducing prototype throughput and increasing overall system costs. Additionally, mechanisms of microfluidic control have been a focal point, while fluid storage prior to manipulation as well as system-level molecular diagnostics demonstrations have remained comparatively underreported.

In this study, we present a low-cost, easy to manufacture microfluidics-based cartridge well suited for molecular diagnostics platforms. The cartridge is able to store buffers and reagents in chambers separated by a custom pressure-driven check valve. Acrylic cartridge components are fabricated using a benchtop laser cutter and micromilling machine, the check valve is made in a custom acrylic mold, and the cartridge is assembled and thermally bonded in a diffusion welder. In this way, high resolution features possible through micromachining and high-throughput fabrication enabled by laser cutting are used in combination with a low-cost bonding method to demonstrate multilayer thermoplastic microfluidic cartridges adequate for prototyping and low-resource environments.

## 2. Concept and Theory of Operation

As the cartridge is intended to be used in low-resource environments, several requirements are inherent to the design. First, a normally closed check valve is required to allow storage of buffers and reagents without premature mixing. Second, low-power and low-voltage operation of the valve is needed for device portability. By utilizing a pressure-driven microvalve, no in-cartridge power is required for valve operation, and low-power activation mechanisms can be held outside of the cartridge. Third, fabrication techniques selected must allow both rapid prototypes and low-cost production without the need for large or expensive machinery. By using a combination of benchtop laser cutting and micromachining, both fast and high-resolution fabrication can be achieved. Lastly, materials selected must be low cost, readily available and suitable for optically based molecular diagnostics platforms. This is accomplished by using readily available optically clear acrylic sheets.

As shown in **Figure 1**, the fabricated cartridge is intended to be pre-filled with a lysing buffer (Chamber 1) and reagents required for Loop-Mediated Isothermal Amplification (LAMP) (Chamber 2). A custom pressure-driven check valve prevents lysing buffer from entering the reagent chamber prematurely, either during storage or sample introduction. After connecting the sample to the cartridge using a press-fit syringe needle, a vacuum pulls the sample into Chamber 1 through an air trap, which prevents any potential sample backfill into the vacuum tubing. The sample is then given time to lyse. After the lysing stage, a vacuum is then pulled via air trap through Chamber 2, with sufficient pressure to open the custom valve and allow the sample / lysing buffer mixture to enter the reagent-based chamber. At this point, appropriate heating and detection mechanisms can be activated for the intended diagnostic method.

**Figure 1.**
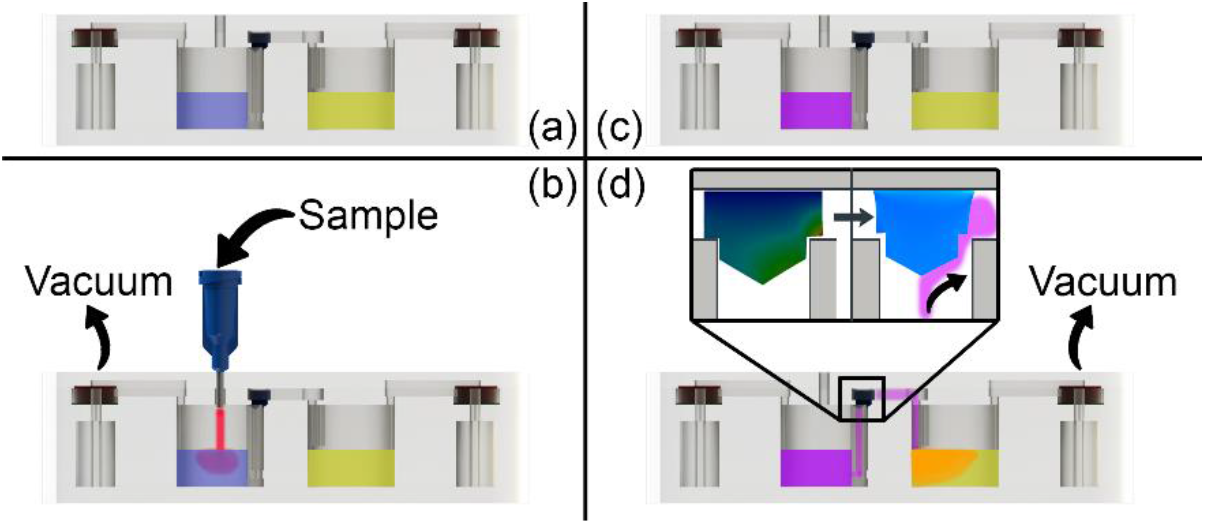
Cartridge theory of operation. a) Two chambers are preloaded with lysing buffer and reagents; b) A sample is drawn into the lysing chamber using a vacuum through an air trap; c) The sample is lysed; d) The sample is pulled with a vacuum past a pressure-driven check valve into the reagent chamber.

## 3. Cartridge Fabrication

Fabrication of the cartridge has been broken down into three major parts: fabrication of the custom check valve, fabrication of the cartridge acrylic components, and overall cartridge assembly via thermal diffusion bonding. Cartridge fabrication steps are shown in **Figure 3**.

### 3.1 Fabrication of check valve

As shown in **Figure 2**, To fabricate the polydimethylsiloxane (PDMS) check valve, a custom mold was created using a standard CO_2_ laser cutter (Universal Laser Systems VLS2.30) and a benchtop Computer-Numerical-Control (CNC) mill (Roland Model A MDX-50). The base of the check valve was created using a 1/32” square endmill, while the pointed plug shape was created with a 1.2mm drill bit. After creating the mold, PDMS (Sylgard 184) was mixed at a 10:1 base-to-curing agent ratio by weight and pulled into a 3mL syringe. The syringe was then used to individually fill each cavity of the mold. The filled mold was then placed under vacuum for 20 minutes, allowing any air bubbles to release from the PDMS. Next, a strip of standard Scotch-brand tape was placed on the mold and pressed to spread out any excess PDMS. The filled mold was then baked at 60°C for a minimum of 4 hours to allow the PDMS to cure. After cooling, the tape was peeled off of the mold, providing multiple check valves which can be easily removed from the tape for use.

**Figure 2.**
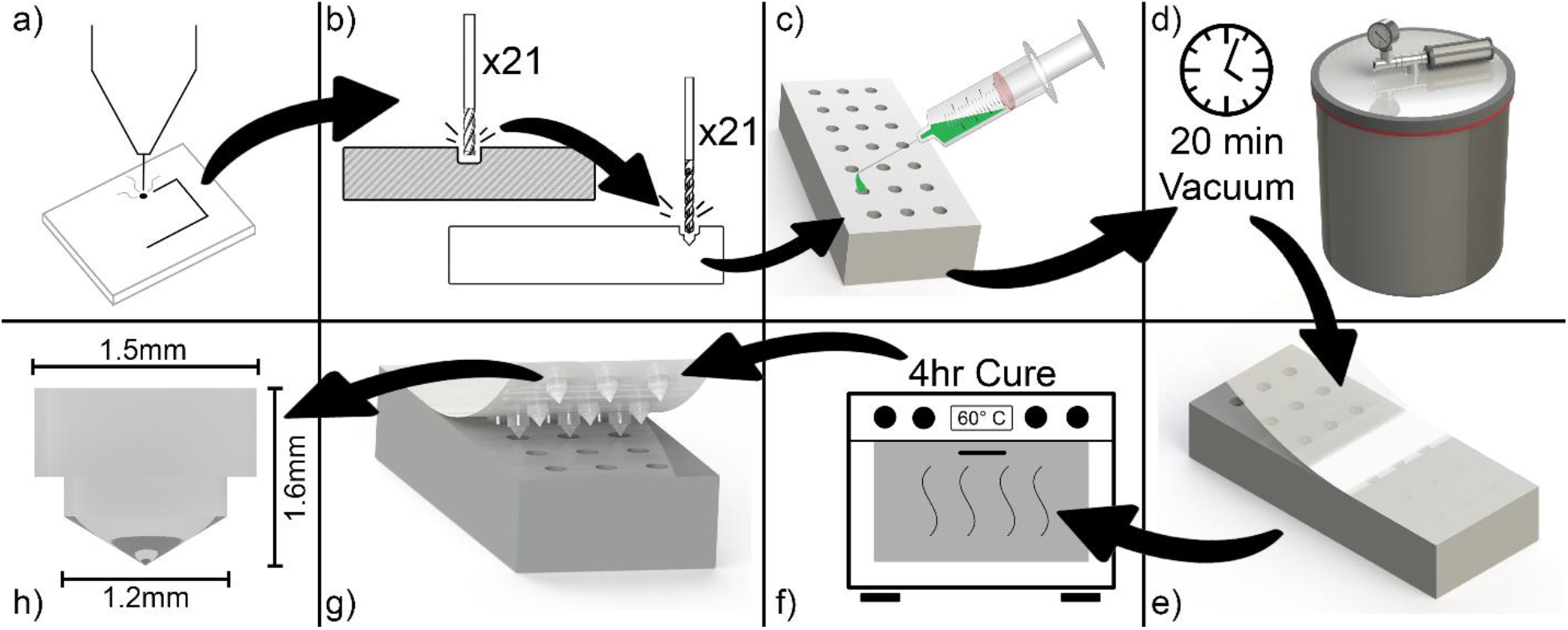
Fabrication of pressure-driven check valve. a) Laser cutting of mold outline; b) Cutting mold using a square and pointed endmill on a benchtop CNC; c) Filling mold with Sylgard 184 PDMS using a 3mL syringe; d) Vacuum-setting mold to get rid of trapped air in PDMS; e) Applying tape layer to mold; f) Mold is cured at 60°F for 4 hours; g) Peeling back tape layer reveals cured check valves; h) Demonstrative dimensions of completed check valve.

**Figure 3.**
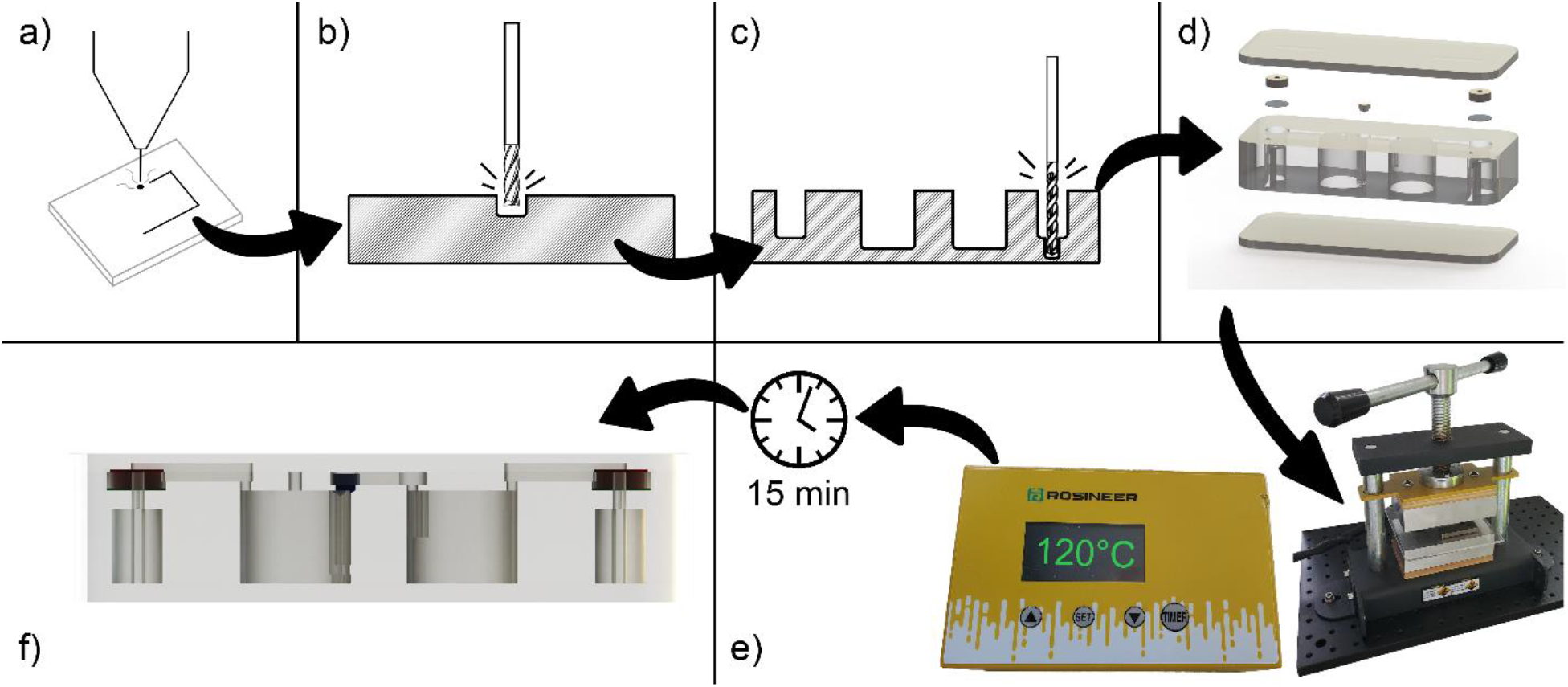
Microfluidic cartridge fabrication. a) Cartridge outline laser cut; b) Milling of chambers and microfluidic channels using a benchtop CNC; c) Drilling of sample input and vacuum connection points performed by the benchtop CNC; d) Assembly of three acrylic pieces, two 0.22um PTFE filters, two gaskets to compress filters in place, and check microvalve; e) Diffusion bonding of assembled components performed at 120°C for 15 minutes; f) Completed microfluidic cartridge containing air traps with filters, two reaction chambers, inlet/outlet ports for vacuum activation of fluid and input of sample, and channels for fluid movement within the cartridge.

### 3.2 Fabrication of acrylic cartridge components

The acrylic cartridge components are fabricated using the standard laser cutter and benchtop CNC described in Section 3.1. To fabricate the top layer, the laser cutter is used to first vector cut the outside edge in 1mm thick acrylic, then raster etch channels which serve as openings between air traps and fluid-holding chambers. Next, the bottom layer outline is vector cut out of the 1mm acrylic sheet. Lastly, 5.6mm thick acrylic is used for the inner layer of the cartridge. First, the outside edge is vector-cut with the laser cutter out of the acrylic. The workpiece is then transferred to the CNC micromilling machine, where a cutout has been created in a sacrificial layer to fit the workpiece, preventing movement during cutting. A single layer of 0.006” thick double-sided adhesive is placed on one side of the pre-cut middle layer, as is standard process for this CNC machine, and the piece is placed in the cutout on the sacrificial layer. The inlet/outlet ports, chambers, and internal fluid flow tunnels are then cut using the CNC with a 1.7mm drill bit, 1/32” square endmill, and 0.7mm drill bit, respectively. Following this operation, the three pieces of acrylic used to create the cartridge are ready for assembly.

### 3.3 Cartridge assembly

Prior to assembling the cartridge, a 0.22µm pore size polytetrafluoroethylene (PTFE) filtration sheet (EZ Flow Hydrophobic PTFE Membrane) is cut using a 3mm diameter hole punch to create filters for the two air traps. The filtration pieces are then placed in their respective places within the middle layer, followed by two rubber gaskets (McMaster-Carr 1182N001) used to compress the filters, holding them in place. The check valve is then placed at the outlet port of the first chamber, pointed-side down.

### 3.4 Diffusion bonding

After assembling the cartridge middle layer subassembly, the top and bottom acrylic layers are added to form a single cartridge. After assembling the three acrylic components, the outer edges are spot-welded using a soldering iron in several places to ensure they remain in place during the diffusion bonding process. Next, the assembly is placed in the thermal diffusion welder (Rosiner GRIP Heat Press) and surrounded with aluminum bars to ensure proper heat transfer from the diffusion welder to the assembly. The assembly is then compressed by manually tightening the lead screw and treated at 130°C for 15 minutes. Lastly, the cartridge is removed from the diffusion welder and allowed to cool to room temperature.

## 4. Experimental characterization of pressure-driven check valve

Following simulation of the check valve, pressure requirements for valve operation were validated as shown in **Figure 4**. First, a colored dye was placed in the sample syringe and connected to the lysing chamber through a press-fit syringe needle. A pressure sensor (Honeywell HSCDRRN001BASA3) communicating to a Teensy 4.1 microcontroller over a Serial Port Interface (SPI) was connected in parallel to the syringe. A vacuum was then slowly pulled through the lysing chamber through the first air trap manually using the syringe. The increasing vacuum pressure was then recorded on a computer through the Teensy microcontroller. Once the dye was observed entering the first chamber, the pressure level was recorded. Following a similar procedure, a vacuum was then pulled through the reaction chamber manually with a syringe connected to the second air trap. The vacuum pressure required to overcome the check valve was then recorded. This process was repeated on 4 cartridges, twice each, to observe any variation in pressure requirements.

**Figure 4.**
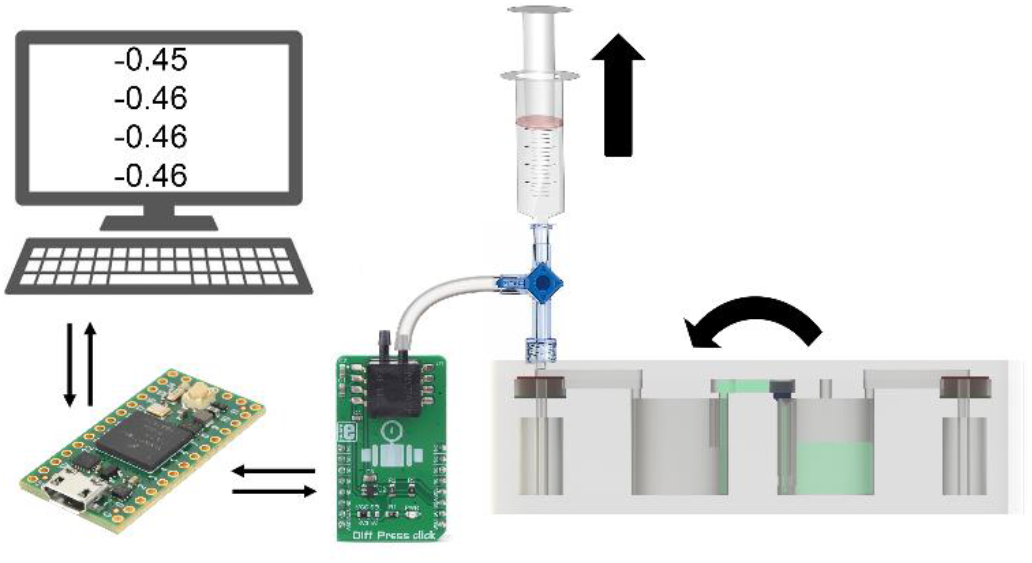
Test setup for valve characterization. Vacuum pulled through external syringe is measured by a differential pressure sensor connected in parallel with the syringe. The pressure sensor displays recorded values on a computer through a Teensy microcontroller.

## 5. Proof of concept validation in custom LAMP genetic testing system

To validate the cartridge application in a molecular diagnostics platform, a simple validation was performed on a custom benchtop LAMP genetic testing system shown in **Figure 5**.

**Figure 5.**
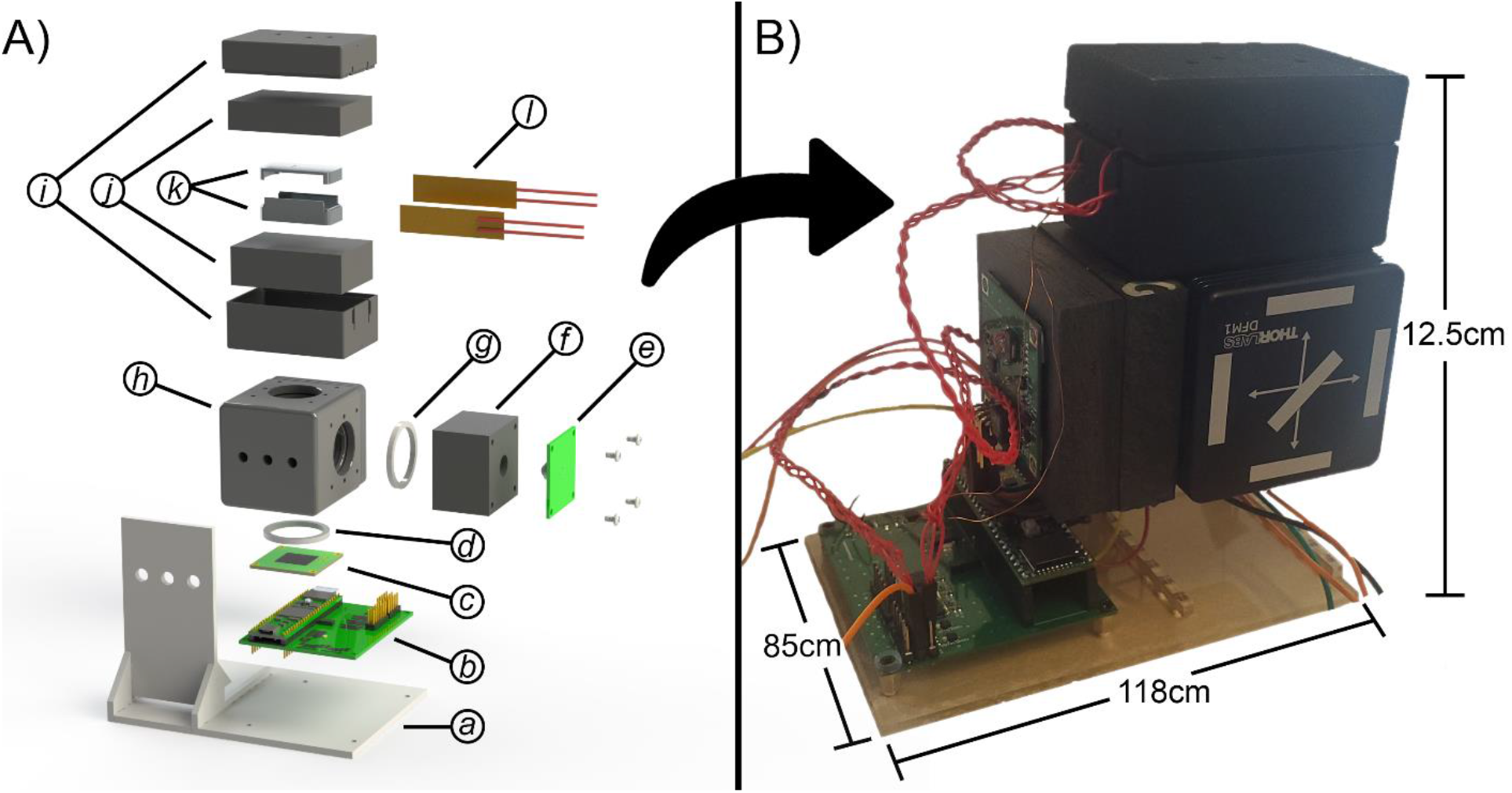
Custom benchtop LAMP system used to validate cartridge for molecular diagnostics platforms. A) Exploded view of the benchtop LAMP system with mounting hardware omitted, showing: a) Custom laser-cut platform to hold the optical, heating, and microcontroller subsystems; b) Teensy 4.1 microcontroller breakout board mounted to custom thermal control circuit board; c) Custom optical collector circuit board; d) Ring gasket applied between optical collector PCB and beamsplitter; e) Custom optical emitter circuit board; f) Custom collimator for optical emitter PCB; g) Ring gasket applied between collimator and beamsplitter; h) Thorlabs beamsplitter; i) Custom 3D-printed heating system enclosure; j) Cut Styrofoam insulation layer; k) Custom aluminum block to house cartridge; l) Thin film heating elements which are mounted to the aluminum block (k); B) Assembled view of benchtop LAMP system used to validate the cartridge for molecular diagnostics platforms.

### 5.1 Benchtop LAMP System

While the focus of the current work is not on the custom LAMP system, a high-level overview of the system is provided for clarity. Briefly, the system is run on a Teensy 4.1 microcontroller breakout circuit board which operates the heating, optical emission, and optical collection systems. The optical emission and collection systems contain custom designed and fabricated circuit boards which are mounted directly to a Thorlabs beamsplitter (DFM-1). Ring gaskets were added between the optical system circuit boards and the beamsplitter to prevent ambient light intrusion from inconsistencies along the circuit board face. Due to the wide beam angle of the lensed LED, a custom collimator was added between the optical emission circuit board and the beamsplitter. The heating system contains a custom CNC-milled aluminum block housed inside of a cut Styrofoam insulation layer to prevent thermal losses to ambient, both of which are housed within a custom 3D-printed enclosure which is mounted to the beamsplitter. Two 25-watt/in^2^ thin-film thermoelectric heaters (Minco HAP6945) are mounted to the aluminum block along with three thermocouples for thermal system feedback. A custom circuit board interfaces the thermocouples and heating elements to the Teensy microcontroller.

In this demonstration, external vacuums are applied to the cartridge via press-fit needle hookups attached to syringes through standard surgical tubing. Upon applying the second vacuum to pull the sample into the reaction chamber, the cartridge is placed in the assembly and heating elements are activated to heat the custom aluminum block (and with it, the cartridge) to 65°C. Thermocouples provide feedback to the system to stabilize at 65°C via pulse-width-modulated power cycling of the heating elements through FETs. From this point, every second the optical emitter LED is activated and the photodetector monitors activity within Chamber 2. Photodetector readings are stored to on-board memory for post-experiment curve analysis.

### 5.2 Testing Protocol

LAMP protocols for testing were adapted from New England Biolabs (NEB) WarmStart LAMP Kit Protocol [30]. Briefly, 12.5µL LAMP warmstart master mix, 0.5µL 50x fluorescent dye, 2.5µL 10x primer mix, 8.5µL de-ionized H_2_O, and 1µL COVID target RNA purchased from NEB were mixed per positive test to create a sample concentration of approximately 100 copies per µL, with the target RNA replaced with de-ionized H_2_O for the negative control test. For multiple simultaneous tests, mixing was performed at once by multiplying required volumes by the number of tests to be performed, then sampling 50µL from the larger volume. As a standard for comparison, a commercially available benchtop LAMP system (MyGo S Real Time Thermal Cycler) was used with standard 100µL PCR tubes. Approximately 50µL of positive sample was manually injected into the custom cartridge using a 1mL syringe.

## 6. Results

### 6.1 Microfluidic Manipulation

After fabricating four cartridges, two trials were performed on each to determine pressure required to introduce the sample and overcome the check valve, as well as any variability in pressure required between trials. First, to pull a sample into the lysing chamber, 1.5 PSI (std. dev. 0.5PSI) was required to fill the chamber in less than 2 seconds. Next, to pull the lysed sample from the first chamber into the reagent chamber, a pressure of 4.5 PSI (std. dev. 0.4PSI) was required. The standard deviation observed may be explained by tolerances in manufacturing techniques and settling variance of the PDMS during the curing period leading to slightly different geometries in the seated section.

### 6.2 LAMP Proof-of-Concept

Following the protocols described above, two positive samples at a concentration of 100 RNA copies/µL and one negative control were created to test the custom cartridge against the commercially available benchtop LAMP system. One positive sample and the negative control were placed in standard PCR tubes in the commercial system, and approximately 50µL of positive sample was injected directly in the cartridge reagent chamber, bypassing the lysing step as the sample mixed contained the target RNA. Following a full trial in each system, data was exported for analysis. Results indicating the characteristic S-curve are shown in **Figure 6**.

**Figure 6.**
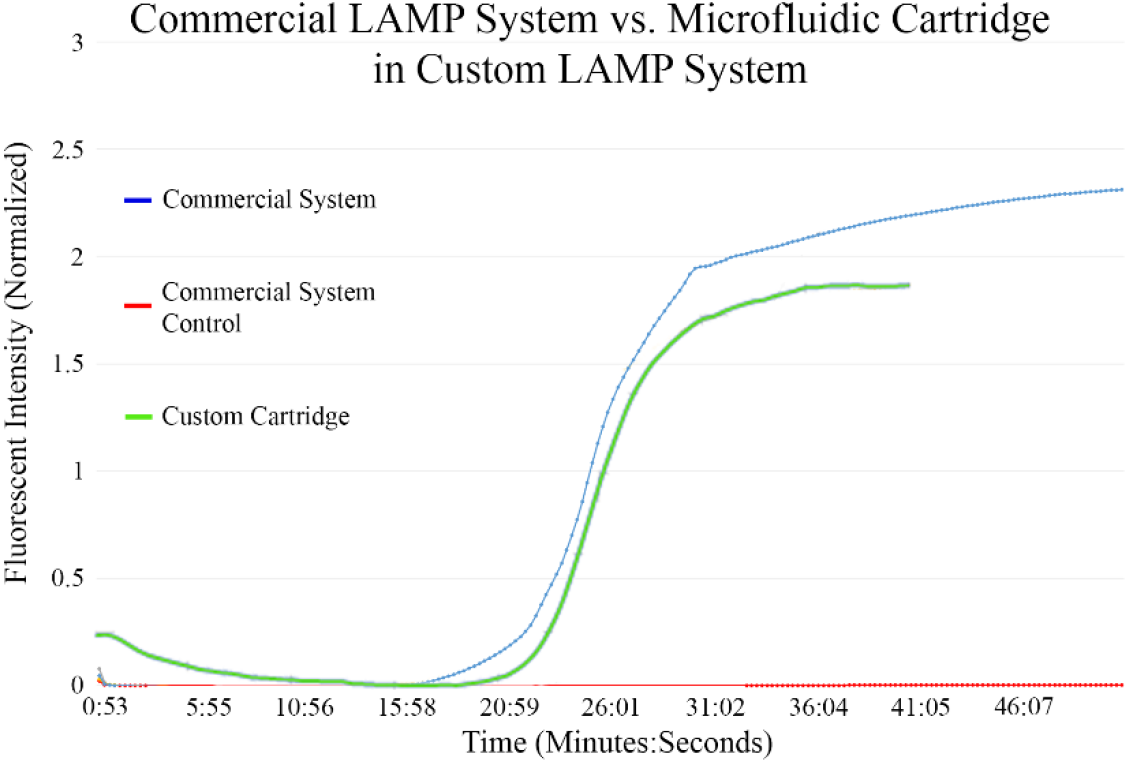
LAMP results. Comparison of fluorescing LAMP results between trials performed on a commercially available benchtop LAMP system and in the custom microfluidic cartridge using a custom benchtop LAMP system. Positive samples were tested at a concentration of 100 copies/µL.

## 7. Conclusion

This work demonstrates the design and fabrication of a thermoplastic microfluidic cartridge incorporating a custom pressure-driven check valve. By combining benefits provided through commonly used laser cutters and less-commonly used benchtop micromachines, the design may be readily fabricated using acrylic and room-temperature curing silicone. Final assembly of the multi-layer microfluidic system was performed on a benchtop thermal diffusion bonding system. The check valve experimentally operated as expected, allowing a sample to be pulled into the first chamber then the second chamber through successive vacuums. Thus, leak-free microfluidic control between two chambers, which may be used for a lysing buffer and reagent storage in molecular diagnostics platforms, was achieved. Initial compressed height and overall height of the valve design may be altered in future designs to adjust the vacuum pressure required to open the valve and allow fluid transfer between chambers. Lastly, a simple LAMP experiment was performed to demonstrate cartridge application in a molecular diagnostics platform.

Overall fabrication time of cartridges is critical to efficient iterative design practices. For a single cartridge, a majority of fabrication time is roughly divided between the thermal diffusion bonding and micromachining steps. Assuming multiple cartridges are to be made simultaneously, relative time per cartridge spent in the diffusion bonding step is reduced as more assemblies may be bonded simultaneously, while the micromachining time remains constant for each cartridge. In other words, overall time diffusion bonding remains constant, while micromachining requires linearly increased time for every cartridge to be fabricated. This machining time may be reduced with higher-powered benchtop machines capable of higher spindle speeds, and thus faster feed rates. Additionally, if speed of fabrication is critical, the benchtop laser cutter may be used to raster-cut the rough-pass of the cartridge, which can be transferred to the micromachining system for a finishing pass.

The unique combination of benchtop laser-cutting and micromachining allows rapid development and testing of microfluidic-based cartridges, as demonstrated with the custom benchtop molecular diagnostics platform. The presented cartridge fabrication method allowing an integrated valve could be widely utilized in both research laboratories and low resource environments to quickly prototype or manufacture devices.

